# Structure-based discovery and *in vitro* validation of selective inhibitors of Chloride Intracellular Channel 4 protein

**DOI:** 10.1101/2022.04.21.489122

**Authors:** Fisayo Olotu, Encarnacion Medina-Carmona, Abdelaziz El-Hamdaoui, Özlem Tastan Bishop, Jose L. Ortega-Roldan, Vahitha B. Abdul-Salam

**Affiliations:** Research Unit in Bioinformatics (RUBi), Department of Biochemistry and Microbiology, Rhodes University, Makhanda, 6139, South Africa; School of Biosciences. University of Kent. CT2 7NJ. Canterbury. United Kingdom; Departamento de Quimica-Fisica. Facultad de Ciencias. Universidad de Granada; Centre for Cardiovascular Medicine and Device Innovation, William Harvey Research Institute, Barts and The London School of Medicine and Dentistry, Queen Mary University of London

**Keywords:** Chloride intracellular channel protein 4, GSH-like catalytic site, Structure-based drug discovery, Computational high-throughput screening, Nuclear magnetic resonance, Allosteric inhibition

## Abstract

Chloride Intracellular Channel Protein 4 (CLIC4) is a novel class of intracellular ion channel highly implicated in tumour and vascular biology. It regulates cell proliferation, apoptosis and angiogenesis; and is involved in multiple pathologic signaling pathways. Absence of specific inhibitors impedes its advancement to translational research. Here, we integrate structural bioinformatics and experimental research approach for the discovery and validation of small-molecule inhibitors of CLIC4. High-affinity allosteric binders were identified from a library of 1615 Food and Drug Administration (FDA)-approved drugs via a high-performance computing-powered blind-docking approach, resulting in the selection of amphotericin B and rapamycin. NMR assays confirmed the binding of the drugs. Both drugs reversed stress-induced membrane translocation of CLIC4 and inhibited endothelial cell migration. Structural and dynamics simulation studies further revealed that the inhibitory mechanisms of these compounds were hinged on the allosteric modulation of the catalytic glutathione (GSH)-like site loop and the extended catalytic β loop which may elicit interference with the catalytic activities of CLIC4. Structure-based insights from this study provide the basis for the selective targeting of CLIC4 to treat the associated pathologies.

## 1. INTRODUCTION

Despite ion channel drugs being almost 18% of currently marketed medications with an estimated global sale of £10 billion, the development of newer drugs, especially for chloride channels, is still lagging. This is mainly due to the lack of efficient pharmacological pipelines targeting these proteins and an incomplete understanding of their precise mechanisms in biological systems^1^. Chloride intracellular channel 4 (CLIC4) belongs to the highly conserved six-membered family of globular proteins (CLIC1-6) which structurally relate to the omega-class glutathione S-transferases (GSTΩs)^2^. Functionally, CLIC proteins are not conventional chloride channels and as well do not function similarly to GST proteins, hence, the majority of their activities do not depend on their roles as channel proteins^2,3^. CLICs are globular proteins and have been associated to varieties of multiorganelle/cellular processes, some of which include tubulogenesis, membrane remodeling, endosomal trafficking, vacuole formation, and cell adhesion ^3–6^. Particularly, CLIC4 is homogenously distributed in the cytosol and exhibits glutaredoxin-like glutathione-dependent oxidoreductase enzyme activity^7^. In the presence of activating molecules (agonists), CLIC4 translocates rapidly and reversibly from the cytosol to the plasma membrane, with the involvement of G-actin-binding protein profilin 1 and actin polymerization induced by Rho and mammalian Diaphanous (mDia) 2 (mDia2) formin^8,9^. Several studies have reported the dynamic association of CLIC4 with effector proteins in the cytosol as well as in the plasma membrane. CLIC4 has been shown to localize in lipid rafts where it interacts with ezrin-radixin-moesin (ERM) proteins that connects receptor proteins in the membrane with submembrane actin cytoskeleton ^10^. Its association with β_1_ integrin also accounts for its modulatory roles in cell adhesion, and migration ^11^. Furthermore, CLIC4 reportedly functions as a scaffold for protein kinases and phosphatases and therefore plays crucial role in the phosphorylation of signaling proteins such CDK2 and CDK6, Smad2/3, p38, IKKβ, MKK6, JNK and SEK1^3,12^. CLIC4 localization and modulatory activities have been reported in the lungs^13^ and bone marrow ^14^ as well as in several cellular types and intracellular organelles^2^ which could explain its significance in multiple cellular and physiological processes.

As a result of its metamorphic roles, CLIC4 has been implicated in various pathophysiological pathways ^15^. For instance, it is well-characterized for its roles in cancer^12,15,16^ and pulmonary arterial hypertension (PAH) ^17,18^. Studies have revealed that CLIC4 regulates multiple stages of angiogenesis and acts upstream of hypoxia-inducible factor (HIF-1α) and vascular endothelial growth factor (VEGF) signaling. This could explain its increased expression in many cancers presenting it as a target for the development of novel cancer therapeutics ^16,19,20^. Suppressing CLIC4 has been shown to decrease cell proliferation, capillary network formation, capillary-like sprouting, and lumen formation ^21,22^. Also, inhibiting CLIC4 correlatively enhanced the phosphorylation of Bcl-2, Bcl-xL and Bad which in turn increased β-cell survival and cytokine-mediated apoptotic resistance ^23^. Similarly, reducing *clic4* expression and blocking downstream interactions may provide a novel way to prevent diabetes-related β-cell apoptosis. More so, CLIC4 was highly expressed in the lungs of patients with pulmonary arterial hypertension (PAH) as compared to healthy patients ^13^ and regulated the activities of the transcription factors, NF-κB and HIF-1, that are responsible for endothelial responses to inflammatory and angiogenic stimuli.

Several attempts have been made to achieve and translate the pharmacological inhibition of CLIC4 for disease treatment but have been unsuccessful to date. This, among many others may be due to its high structural and sequence similarity with other CLIC proteins as well as the GSTΩs ^7,24^. Small-molecule inhibition of CLIC proteins has been commonly achieved with the use of indanyloxyacetic acid-94 (IAA-94), an intracellular chloride channel blocker designed based on the GST inhibitor, ethacrynic acid ^25^. This compound however lacks specificity and binds with various members of the CLIC family (SspA, CLIC1, CLIC3, CLIC4, and CLIC5) ^16,26,27^ and other chloride channels family of proteins such as pannexin1^28^, bestrophin^29^, Calcium-activated chloride channels, Volume regulated anion channels (VRAC)^30^. Other known chloride channel blockers such as A9C and DIDS have been tested on CLICs where only A9C but not DIDS is known to act on CLICs ^31^. A9C was recently shown to also inhibit the enzymatic activity of CLICs ^7^. Pharmacological inhibition of CLICs by IAA-94 is shown to reduce tumor growth as well as prevent neurodegeneration ^32^. Moreover, given the functional versatility of the various CLICs and the non-specific activities of IAA94 and A9C, future research is needed to focus on chemical derivatives or new molecules with improved specificity.

The crystal structure availability of the soluble form of CLIC4 provides an advantage that can be exploited using *in silico* structure-based techniques. Therefore, in this study we implemented integrative computer-based and experimental methods to: (i) identify new druggable allosteric sites on CLIC4; (ii) blind-screen and discover hit inhibitor compounds specific for CLIC4; (iii) validate the inhibitory potentials of the hit compounds *in vitro* (iv) investigate CLIC4 allosteric inhibitory mechanisms using GPU-accelerated molecular dynamics (MD) simulations. We expect that findings from this study will contribute significantly to therapeutic interventions in various CLIC4-mediated pathologies.

## 2. COMPUTATIONAL METHODOLOGY

### 2.1. Retrieval of the protein three-dimensional structure and preparation

The three-dimensional (3D) structure of CLIC4 was obtained from the Protein Data Bank (PDB) with entry 2AHE^33^, and was prepared for subsequent analyses on the graphic user interface (GUI) of UCSF Chimera. For comparative modeling and analysis, the 3D structure of the CLIC1 homolog complexed with glutathione (GSH) was also obtained from PDB with entry 1K0N^34^. This was done so as to correctly map out the GSH-binding region which is reportedly highly conserved in all human CLIC homologs, particularly the catalytic cysteine ^5^. System preparation involved the removal of co-crystallized molecules such as crystal waters from both proteins, and the non-GSH binding monomer from CLIC1. More so, both CLICs were structurally superposed using Needleman-Wunsch algorithm (UCSF Chimera MatchMaker) to define the GSH binding region in CLIC4. This implemented a pairwise alignment of the sequences which were then fitted per aligned-residue pair. CLIC4 residues at a distance of 5Å from the crystal GSH were mapped accordingly to constitute the GSH-binding region in CLIC4.

### 2.2. Identification, cross-validation and characterization of potential allosteric sites

Multiple predictive algorithms were implemented to identify possible allosteric sites on CLIC4 asides the GSH-binding region mapped in section ***1.1***. This approach is in line with some previous studies ^35–39^ and important to predict consensus sites (across the algorithms) with high potentials for druggability. Tools employed for allosteric site prediction and characterization in this study include SiteMap^40^, DeepSite ^41^, FTMap ^42^, DogSiteScore ^43^, and ProBIS ^44^. The targetability of allosteric pockets predicted by SiteMap is further based on properties that include surface exposure, hydrophobicity, hydrophilicity, and druggability, as measured by the Halgren’s and DogSite scores ^40^. The consensus allosteric pocket as commonly predicted across the five algorithms was then mapped and characterized based on these intrinsic attributes.

### 2.3. Computational high-throughput screening and hit identification

A library of 1,615 FDA approved compounds was retrieved from the ZINC15 repository (http://zinc15.docking.org/substances/subsets/fda/), with each constituent compounds subjected to structural optimization, protonation and energy minimization using Open Babel and AutoDock tools for the final conversion into .*pdbqt* formats. The prepared ligands were allowed to be flexible and then used to the screen the target protein (rigid) across its entire surface. This is a blind docking approach which allows small-molecule compounds to ‘non-restrictively’ bind to preferred sites on their target proteins based on affinity and interaction complementarity. In addition, this approach is important to further validate the potential druggability of pockets predicted in section ***1.2***. UCSF Chimera-integrated AutoDock Vina was used to calculate the docking coordinates which include box sizes *x, y, z*; 47.73, 53.92, 60.38 and centers *x, y, z*; -2.46, -12.84, 29.82. As a positive control, IAA-94, a widely reported ‘non-selective’ CLIC inhibitor was blindly docked to CLIC4 using the same coordinates, since the exact IAA-94 binding site on CLIC4 has not been clearly defined. The screening experiment was then performed using AutoDock Vina integrated in a high-performance computing (HPC) cluster. The resulting docked posed of the inhibitor compounds with respect to their scores were visualized in PyMOL GUI after which they were filtered based on their binding energies (affinity), non-preferentially to the GSH binding site and ultimately, their pharmacological relevance (usage). Taken together, the top 10 hits with potential allosteric selectivity for CLIC4 were selected for further evaluation.

### 2.4. Molecular dynamics (MD) simulations of protein systems

Following *in vitro* validative experiments for the predicted hits, two inhibitor-protein complexes together with the unbound and IAA-94-bound proteins (controls) were prepared for long-timescale MD simulations. This was carried out on the AMBER18 Graphical Processing Unit (GPU) and its integrated modules. ^45^ The FF14SB forcefield was used to define the protein parameters and antechamber/parmchk modules for ligand parameterization. Likewise, coordinate and topology files for the unbound and inhibitor-bound proteins were defined with the LEAP program. This program, also, was used to add counter Na^+^ and Cl^-^ ions to neutralize and solvate the systems in a TIP3P water box of size 10Å. Partial minimization was first carried out for 5000steps using a 500kcal mol^-1^. Å^2^ restraint potential followed by another 2500 steps of full minimization without restraints. The systems were then heated in a canonical (NVT) ensemble with a 5kcal mol^-1^ Å^2^ harmonic restraints gradually from 0 – 300k for 50ps, followed by a 250000 equilibration steps in an NPT ensemble at constant temperature of 300K without restraints. Atmospheric pressure was maintained at 1bar with a Berendsen barostat ^46^ while each protein system was subjected to a production run of 500ns. Corresponding trajectories were saved at every 1ps time-frame until the end of the simulation and were analysed using CPPTRAJ followed by data plot analyses using Microcal Origin software ^47^ and an in-house R-script. Snapshots were also taken and analyzed to monitor structural events and ligand interaction dynamics across the trajectories on UCSF Chimera user interface (GUI), PyMOL and Discovery Studio Client. ^48^

### 2.5. Calculations of binding free energies and per-residue decomposition

Differential binding affinities of the validated allosteric CLIC4 inhibitors and control compound were evaluated using the Molecular Mechanics/Generalized Born Surface Area (MM/GBSA) method. Binding energy profiles for these compounds and their corresponding energy components were estimated using 1000 snapshots from the terminal 50ns MD trajectories where the systems exhibited conformational stability. This helps minimize the effects of conformational entropy on the energy calculations, which is mathematically expressed as follows:

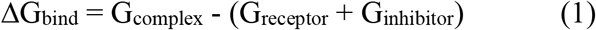

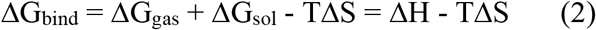

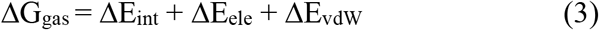

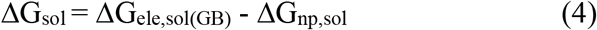

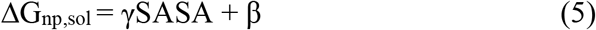

Accordingly, internal (ΔE_int_), electrostatic (ΔE_ele_) and van der Waals (ΔE_vdW_) energies sum up the gas phase energy (ΔG_gas_) while the solvation free energy (ΔG_sol_) is defined by the polar solvation (ΔG_ele,sol_) and non-polar contribution to solvation (ΔG_np,sol_) terms. The MM/GBSA method was used to estimate the Generalized Born (GB) for ΔG_ele,sol_ while linear relationship between the surface tension proportionality constant (γ = 0.0072 mol^-1^ Å^-2^), solvent accessible surface area (SASA, Å^2^), and β constant was used to solve ΔG_np,sol_.

## 3. EXPERIMENTAL VALIDATION METHODS

### 3.1. Protein Expression and Purification

The Human CLIC4 gene was cloned into a pET-28a vector containing an N-terminal hexahistidine tag and a TEV cleavage site. CLIC1 was expressed recombinantly in M9 minimal media supplemented with 15N NH4Cl in the C43 *E.coli* strain (Lucigen). The cells were lysed by sonication, and the membrane and soluble fractions were separated by ultracentrifugation at 117734 g. The soluble fraction was purified separately in the absence of any detergent using affinity chromatography with Ni IMAC. The elutions were pooled and cleaved with TEV protease, and subsequently, gel filtrated using a Superdex200 Increase column (GE) in either 20 mM HEPES buffer with 20 mM NaCl at pH 7.4.

### 3.2. Nuclear Magnetic Resonance (NMR) titrations

The selected high-affinity hit compounds were tested for binding using NMR. Spectra were acquired on a Bruker Avance III spectrometer at a proton frequency of 600MHz using a QCIP cryoprobe. ^15^N TROSY HSQC of 15N labeled CLIC4 were collected in the absence and presence of a two molar excess of each drug or an equivalent volume of DMSO.

### 3.3. Culture of Human Umbilical Vein Endothelial Cells (HUVEC)

HUVECs were purchased from PromoCell and were cultured in endothelial growth medium-2 (Promocell) supplemented with 1% penicillin/streptomycin (Gibco) in 1% gelatin-coated dishes at 37°C under normoxic conditions (20% O2, 5% CO2). The cells were cultured to 70%–80% confluence and were treated with ZnCl_2_ (10 μM), with or without amphotericin B (AMPhB) (10 μM) or rapamycin (RAPA) (10 μM).

### 3.4. Immunocytochemistry analysis of CLIC4 membrane translocation

CLIC4 localization within HUVECs was determined by immunocytochemistry analysis based on a previously described protocol ^17^. Briefly, HUVECs were cultured in 24-wells on gelatin-coated 13mm glass coverslips. After 24 hours, cells were either untreated or treated with 10μMZnCL2 with/without 10 μM RAPA or AMPhB for 8 hours. The cells were then washed with phosphate-buffered saline (PBS), fixed with 4% paraformaldehyde in PBS for 10 min and permeabilised for 10 minutes with 0.3% Triton X-100 in PBS. Non-specific protein-protein interactions were blocked with incubating the cells in 1% BSA for an hour followed by incubation with anti-CLIC4 antibody ((Santa Cruz Biotechnology, clone 356.1, 1/100 dilution) and anti-VE cadherin antibody (Bio-Techne, AF938-SP, 1/500 dilution) overnight at 4 °C. Cells were then washed with PBS and incubated with Alexa Fluor 488 anti-rabbit IgG (green, 1/200 dilution) and Alexa Fluor 488 anti-rabbit IgG (green, 1/200 dilution) for 1 h in the dark. Following a PBS wash step, the cells were mounted with Vectashield Antifade mounting medium with DAPI. All microscopy slides were viewed with a Zeiss LSM-880 confocal microscope using 405 nm, 488 nm, 633 nm lasers. All images were processed with Zen Black and Zen Blue software.

### 3.5. Wound healing assay to validate endothelial cell responses of drugs

HUVECs were grown to confluency in a 6-well plate and a micro pipettor was used to generate a 1-mm wide scratch on the bottom of the 6-well plate. Cells were then gently washed with PBS and were either untreated or treated with VEGF (ug/mL) with/without the drugs in reduced serum media (OptiMem) for 18 hours. Microscopy was used to observe and photograph cell migration to the scratch area and estimate the effects of drugs on wound healing.

## 4. RESULTS

### 4.1. Combinatorial search algorithms identified two de novo and druggable allosteric sites

Identifying putative allosteric CLIC4 sites (other than the GSH-binding site) for pharmacological targeting was achieved using integrative method that involves multiple site prediction algorithms; SiteMap^40^, DeepSite ^41^, FTMap ^42^, DogSiteScore ^43^, and ProBIS ^44^. This integrative approach is essential to validate and cross-validate the allosteric and druggable potentials of these sites. Primarily, three sites (Sites 1-3) were predicted by SiteMap and ranked based on their potentials (Figure 1A and B, Table 1). More so, global pocket descriptors were estimated, which include the cavity size, volume, enclosure, hydrophobicity, and hydrophilicity to define the chemical tractability (druggability) as well as the morphology of the predicted sites. Site 1 has the highest propensity for druggability with SiteScore = 0.865 and Dscore = 0.871 (threshold: SiteScore ≥ 0.8; Dscore > 0.83) but corresponds with the known CLIC4 GSH-binding site located at the N-terminal domain (residues 1→90). The second predicted region is a component of the α-helical C-terminal domain (residues 100→253). More specifically, Site 2 lies around the flexible foot loop region (residues 159→175) and with estimated SiteScore and Dscore of 0.840 and 0.797 respectively. Morphologically, Site 2 constitute a flexible loop region that connects the N-terminal domain to the α-helical C-terminal domain. Site 3, which is located adjacent to the GSH binding site has relatively low cavity size (72 A^2^), volume (140.973 A^3^) and hydrophobicity (0.029) which correlates with lower SiteScore and Dscore of 0.710 and 0.671 respectively, although with a tendency for druggability (intermediate), as predicted by the DogSiteScore algorithm.

**Table 1:**
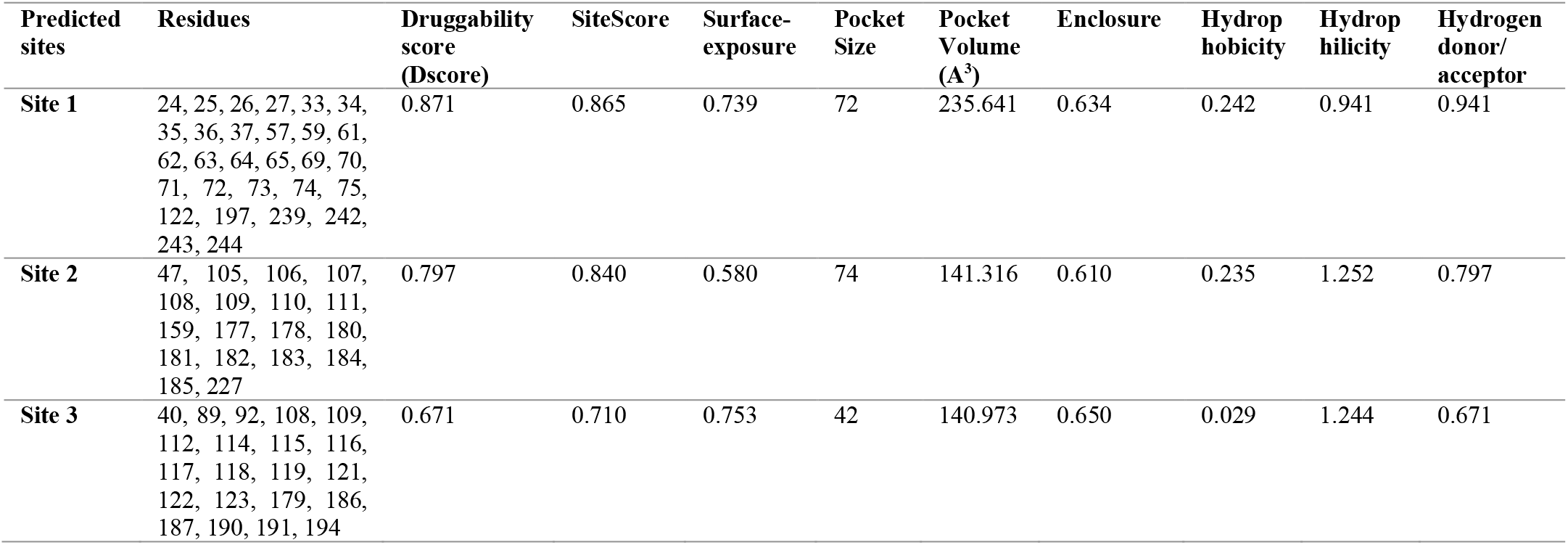
SiteMap identification of potential druggable sites on CLIC4 and characterization

| Predicted sites | Residues | Druggability score (Dscore) | SiteScore | Surface-exposure | Pocket Size | Pocket Volume (Å<sup>3</sup>) | Enclosure | Hydrophobicity | Hydrophilicity | Hydrogen donor/acceptor |
| --- | --- | --- | --- | --- | --- | --- | --- | --- | --- | --- |
| Site 1 | 24, 25, 26, 27, 33, 34, 35, 36, 37, 57, 59, 61, 62, 63, 64, 65, 69, 70, 71, 72, 73, 74, 75, 122, 197, 239, 242, 243, 244 | 0.871 | 0.865 | 0.739 | 72 | 235.641 | 0.634 | 0.242 | 0.941 | 0.941 |
| Site 2 | 47, 105, 106, 107, 108, 109, 110, 111, 159, 177, 178, 180, 181, 182, 183, 184, 185, 227 | 0.797 | 0.840 | 0.580 | 74 | 141.316 | 0.610 | 0.235 | 1.252 | 0.797 |
| Site 3 | 40, 89, 92, 108, 109, 112, 114, 115, 116, 117, 118, 119, 121, 122, 123, 179, 186, 187, 190, 191, 194 | 0.671 | 0.710 | 0.753 | 42 | 140.973 | 0.650 | 0.029 | 1.244 | 0.671 |

**Figure 1:**
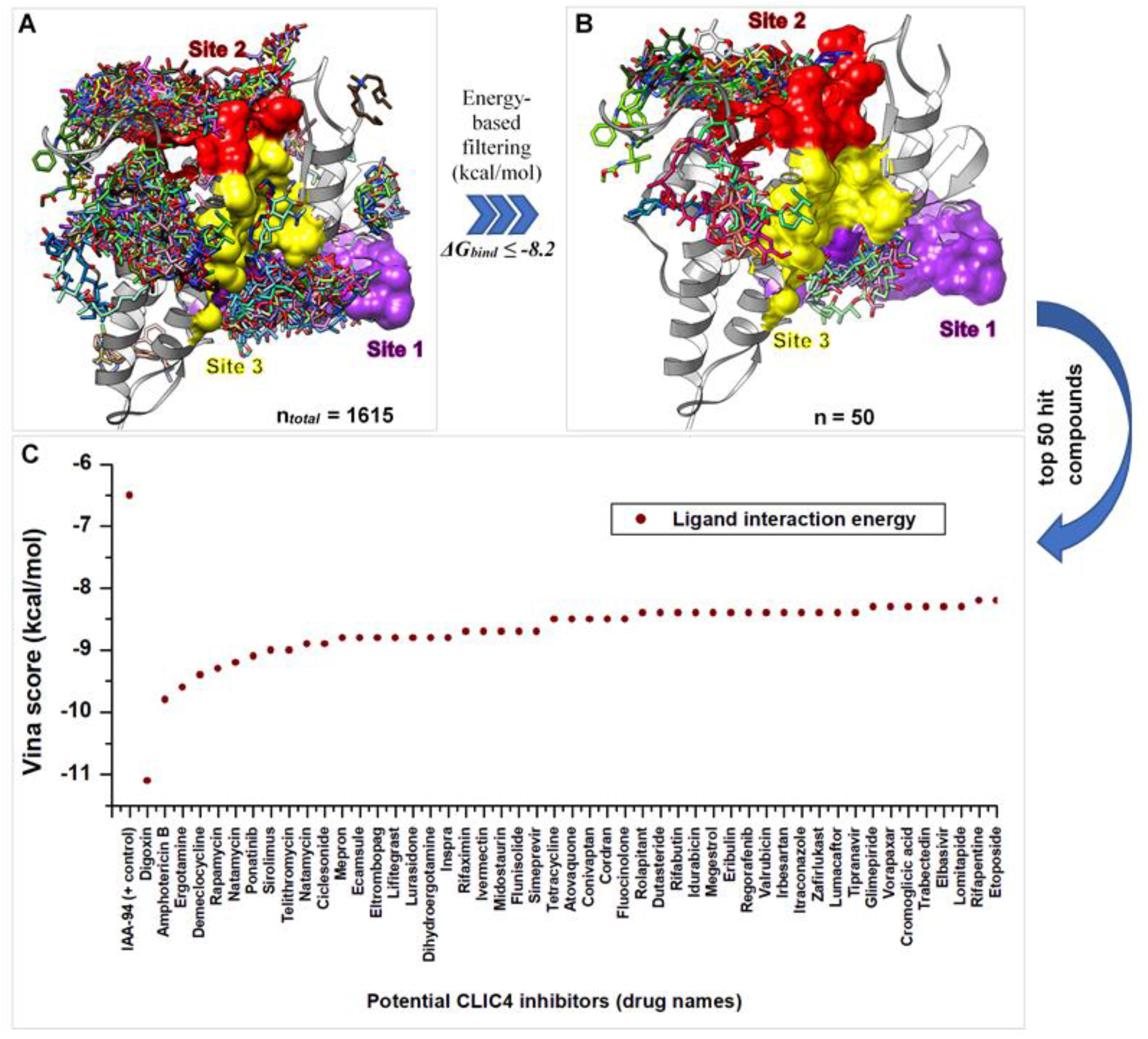
3D pocket localization of the predicted Sites 1-3 and blind-docking/screening and selection of potential hit compounds. **[A]** Blind docking results showing all 1615 FDA compounds binding across various CLIC4 cavities, including the predicted Site 1 (magenta), Site 2 (red) and Site 3 (yellow) all shown in surface representation **[B]** Energy filtering results of the top 50 hit compounds binding to their preferred target sites on CLIC4. [C] Plot showing the interaction energies of the respective compounds in the top 50 set.

As estimated, Site 3 has the highest surface exposure (0.753) as compared to Site 1 (0.739) and Site 2 (0.580) with an enclosure 0.650 for a less well-defined cavity. Functionally, Site 3 residues are located within the α-helical C-terminal domain (residues 100→253) and more specifically proximal to the nuclear localization sequence (NLS, residues 199-206).

Cross-validative predictions to further support the potentials of these predicted pockets were carried out using DogSiteScore, FTMap, DeepSite, and ProBIS (Table 2). Site 1 was commonly predicted across all the four algorithms while two (FTMap and DogSiteScore) complementarily identified residues that constitute Site 2. Moreover, DogSiteScore, PROBIS and DeepSite mapped regions that correlated with the primarily identified Site 3. Druggability estimation (threshold > 0.5) by DogSiteScore further revealed a DrugScore of 0.6 (Site 1), 0.53 (Site 2) and 0.78 (Site 3).

**Table 2:**
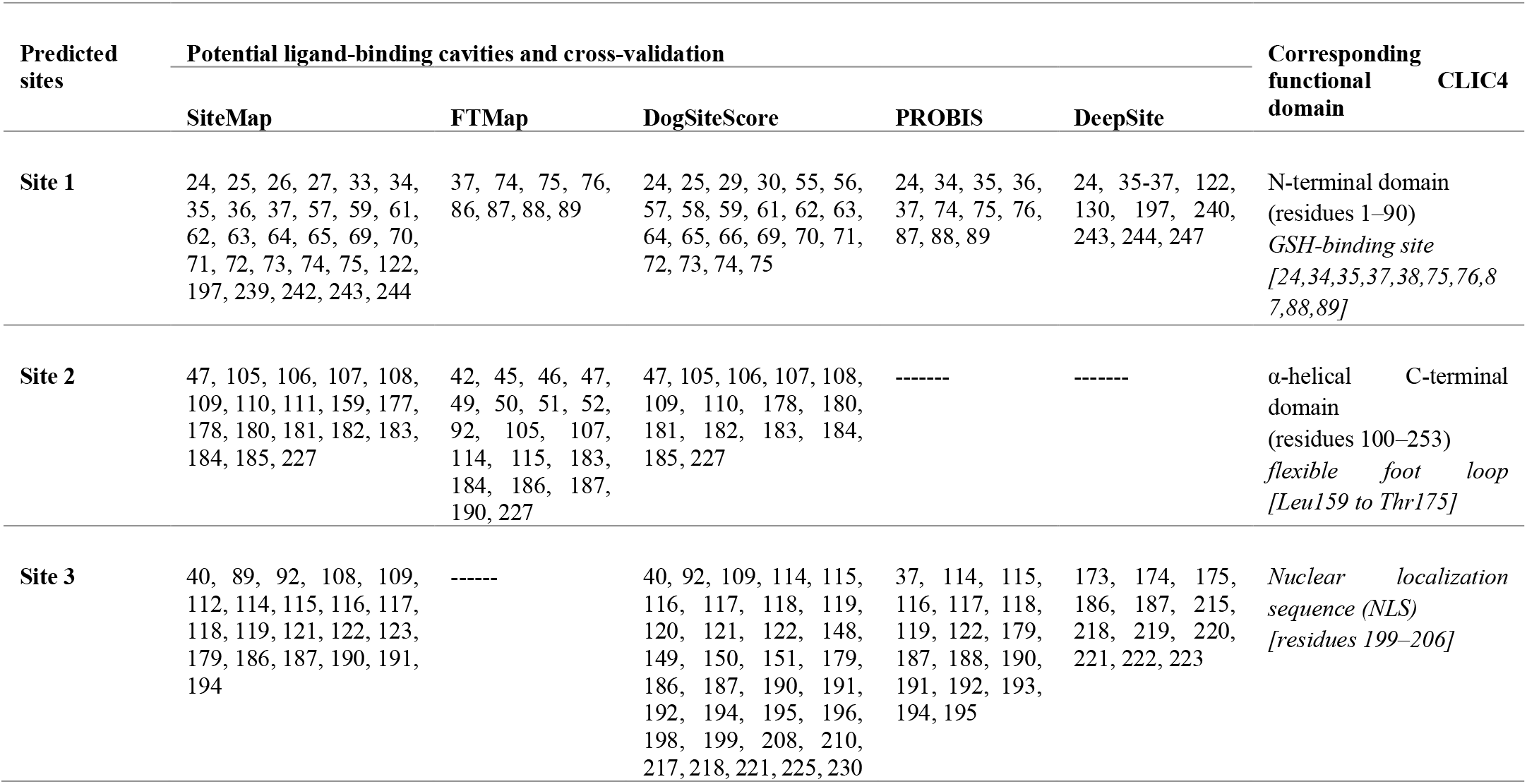
Cross-validation of primarily identified druggable sites using multiple prediction methods

### 4.2. In silico screening and site-directed energy-based sorting led to the selection of site-specific high-affinity hit compounds

This experiment was executed on a HPC-integrated Autodock Vina and entailed a blind screening approach aimed at identifying specific hits (from the 1,615 FDA approved compound library) that bind allosterically to the target protein based on preferentiality.

Results were further analyzed based on agreement with allosteric sites prediction earlier performed. Across all the 1,615 compound set, energy scores ranged from -2.3 kcal/mol (lowest affinity → ZINC8034121: cysteamine) to -11.1 kcal/mol (highest affinity → ZINC242548690: digoxin). An *in-house* filtering algorithm was used to sort compounds with non-GSH site (Site 1) binding activities and interaction energy (*ΔG*_*bind*_) ≤ -8.0 kcal/mol.

Based on energy scoring, results for the top 50 potential CLIC4 inhibitors, including the control compound, IAA-94, were curated and presented in Supplementary Table 1. The screened compounds exhibited high diversities in their inhibitory mechanisms against CLIC4 by binding to different cavities based on their complementary interaction affinity (Figures 1A and B). However, most of the top 50 hit compounds exhibited selective binding to the Site 2 region with variations in their binding energy values (Figure 1). As estimated, the docking score of IAA-94 was -6.4 kcal/mol and it showed a much lower affinity compared to the top 10 hits with scores between - 8.8 to -11.1kcal/mol. The compound with the most inhibitory potential (based on energy scoring) is digoxin ((*ΔG*_*bind*_) = -11.1 kcal/mol) which binds at the predicted Site 2, proximal to the flexible foot loop. Structural (visual) analysis revealed hydrogen bonds with Met184 and Ile163, electrostatic interaction (attractive charge) with Asp161, hydrophobic interactions with Pro158, Leu159, Lys172, Ile163, Lys110, Leu105 and Leu107. Unfavorable interactions however occurred with Arg176 and Glu183. Binding regions and modes for other selected top hits are presented in Table 3 and Figures 2 (0-10). More so, asides AMPhB which binds proximally to the predicted Site 3, other hit compounds (from the top 10 subset) majorly exhibited binding at Site 2 with extended interactions into the flexible foot loop region: Ergotamine binds at the predicted Site 2 and exhibited a binding pattern similar to digoxin by extending into the flexible foot loop.

**Table 3.**
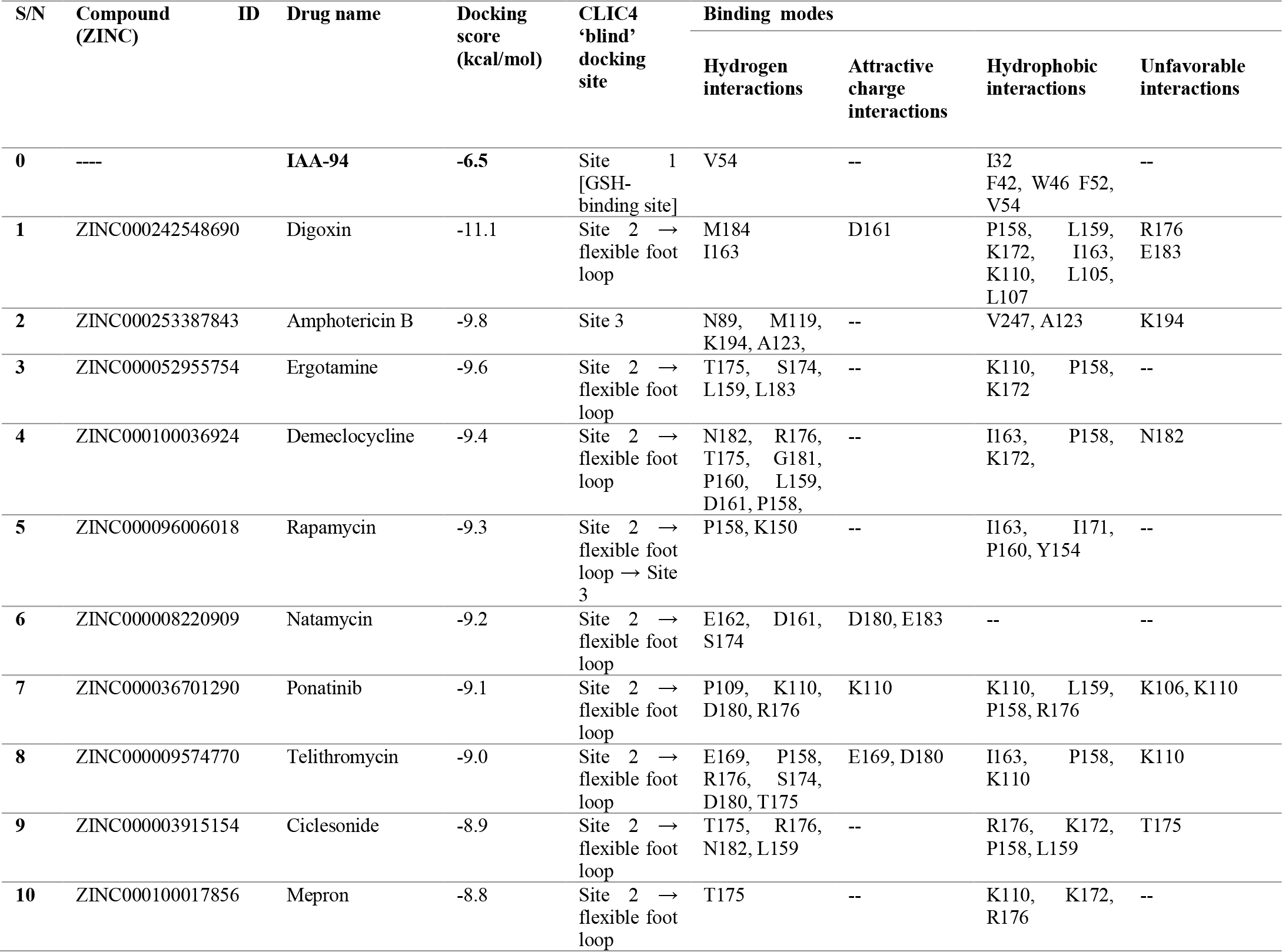
detailing docking results of the top 10 hit compounds

**Figure 2:**
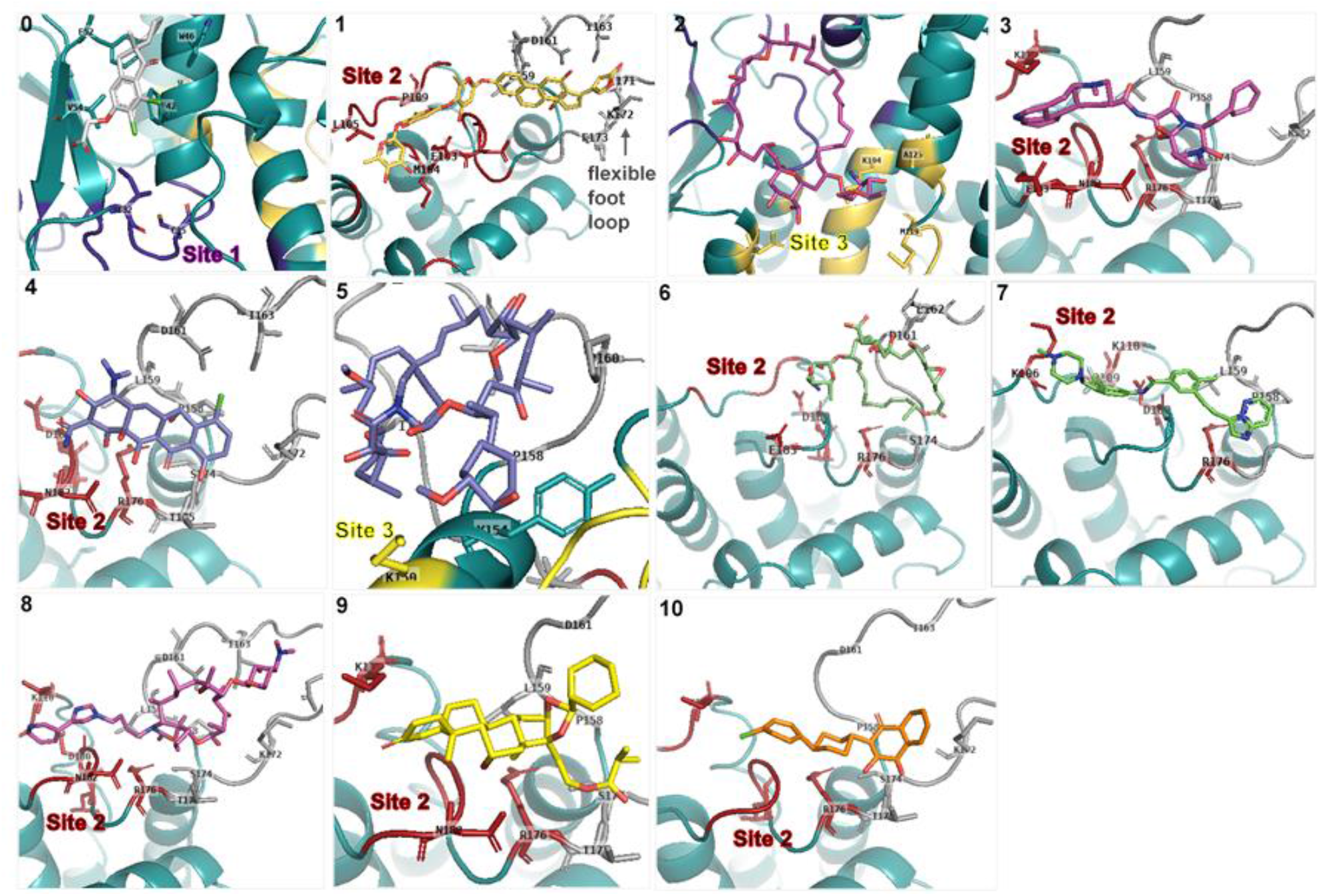
Binding modes and target sites of top 10 potential inhibitors of CLIC4. 0. IAA94 1. Digoxin 2. Amphotericin B 3. Ergotamine 4. Demeclocycline 5. Rapamycin 6. Natamycin 7. Ponatinib 8. Telithromycin 9. Ciclesonide 10. Mepron

Demeclocycline also binds at Site 2 more specifically at the interface of the flexible foot loop. Rapamycin (RAPA) binds around the predicted Site 2, making contact with some residues of the flexible foot loop and Site 3 to bind stably.

Natamycin extends more into the flexible foot loop at the interface with the Site 2 region. Ponatinib binds similar to ergotamine and traverses the Site 2 into the flexible foot loop. Telithromycin binds to the Site 2 region and extends into the flexible foot loop as well, a binding pattern similar for Ciclesonide and Mepron. On the contrary, visualisations revealed that IAA-94 displayed binding around the predicted Site 1 region which corresponds to the GSH binding domain. Hydrogen bonds were observed with Val54 while hydrophobic interactions were observed with Ile32, Phe42, Trp46, Phe52, and Val54.

For experimental testing, AMPhB and RAPA were selected in addition to the control compound, IAA-94, firstly, due to their unique binding positioning among the top 10 compound set. According to our findings, both ligands were uniquely binding away from the catalytic (GSH-binding) site and the predicted Site 2 region (unlike other hits). Secondly, their pharmacological relevance was considered over other compounds in the top 5, particularly with regards to therapeutic usage. For instance, Digoxin is used for treating cardiovascular, however, its long term usage has been associated with incidences of life-threatening conditions like heart attack^49^, hence was not selected for further *in vitro* evaluation.

#### 4.3. AMPhB and RAPA exhibits direct CLIC4 binding and induced significant structural changes

^15^N-^1^H HSQC NMR spectra are widely used to monitor protein-small molecule interaction due to their sensitivity to changes in the chemical environment of individual amino acids. To validate the binding of RAPA and AMPhB to CLIC4, we monitored the changes in ^1^H-^15^N NMR resonances of CLIC4 in the presence and absence of these drugs, and compared it to the effect of the non-selective chloride channel inhibitor IAA-94.

Upon addition of 2 molar excess of AMPhB, significant chemical shift changes were observed for a subset of CLIC4 resonances (Figure 3). RAPA could only be added to an approximate 0.2 molar excess due to its low solubility in aqueous solution, but still induced moderate changes in the NMR spectra. IAA-94, on the other hand, failed to induce any significant chemical shift changes in CLIC4 spectra, indicating that IAA-94 cannot bind to CLIC4 in its soluble state.

**Figure 3:**
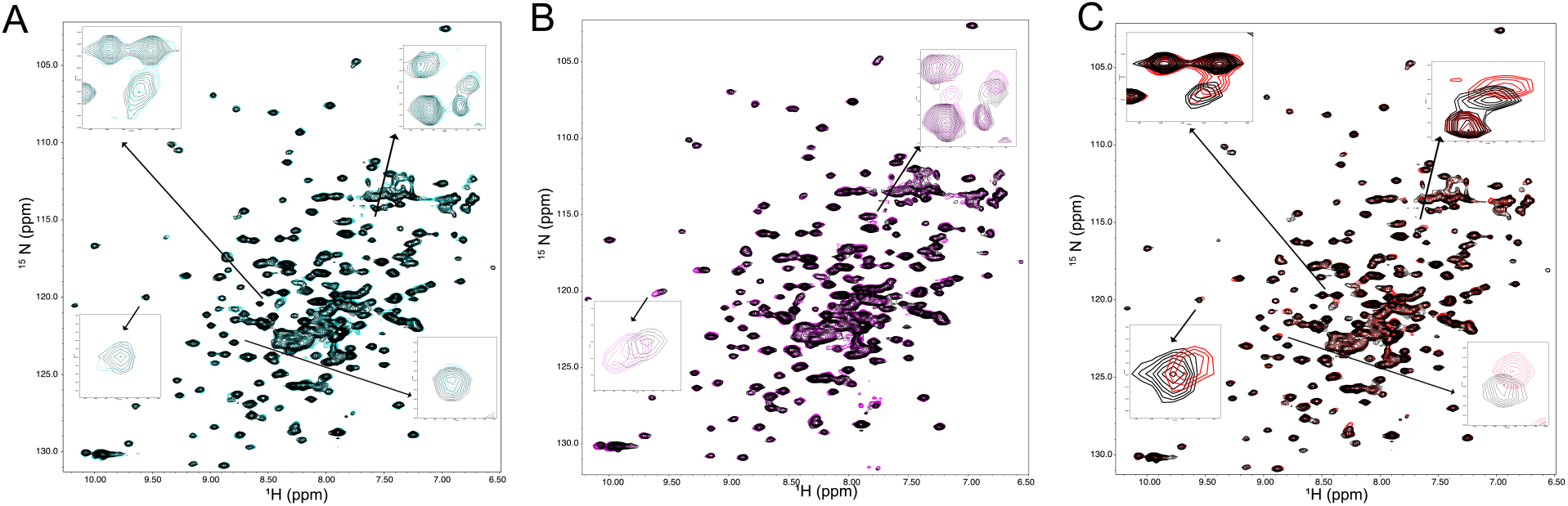
Binding of amphotericin B and rapamycin to CLIC4. ^1^H-^15^N TROSY HSQC spectra of CLIC4 in the absence (black) and presence of two molar excess of IAA-94 (A, cyan), rapamycin (B, magenta) and amphotericin B (C, red). Regions showing significant chemical shift changes upon addition of Rapamycin and Amphotericin B are expanded and compared to similar regions in the IAA-94-bound spectrum.

### 4.4. AMPhB and RAPA inhibits CLIC4 membrane translocation and ameliorates oxidative stress in endothelial cells

Levels of CLIC4 are known to increase under cellular stress conditions such as oxidative stress in a variety of cells ^50^. We have previously shown that oxidative stress promotes significantly higher expression of CLIC4 in HPAECS resulting in endothelial dysfunction ^17^. This was accompanied by deleterious endothelial responses including an increase in VEGF mediated angiogenesis^17^.

Here, we investigated the inhibitory properties of AMPhB and RAPA on the deleterious effects of CLIC4 on HUVECs.

Under normal condition, CLIC4 is modestly expressed or mostly limited to cell cytosol. HUVECs treated with hydrogen peroxide induces oxidative stress and significantly increased CLIC4 levels in all cellular compartments including plasma membrane. These effects were partly reversed by the addition of AMPhB and RAPA, indicating inhibition of CLIC4 response to oxidative stress (Figure 4, top panel).

**Figure 4:**
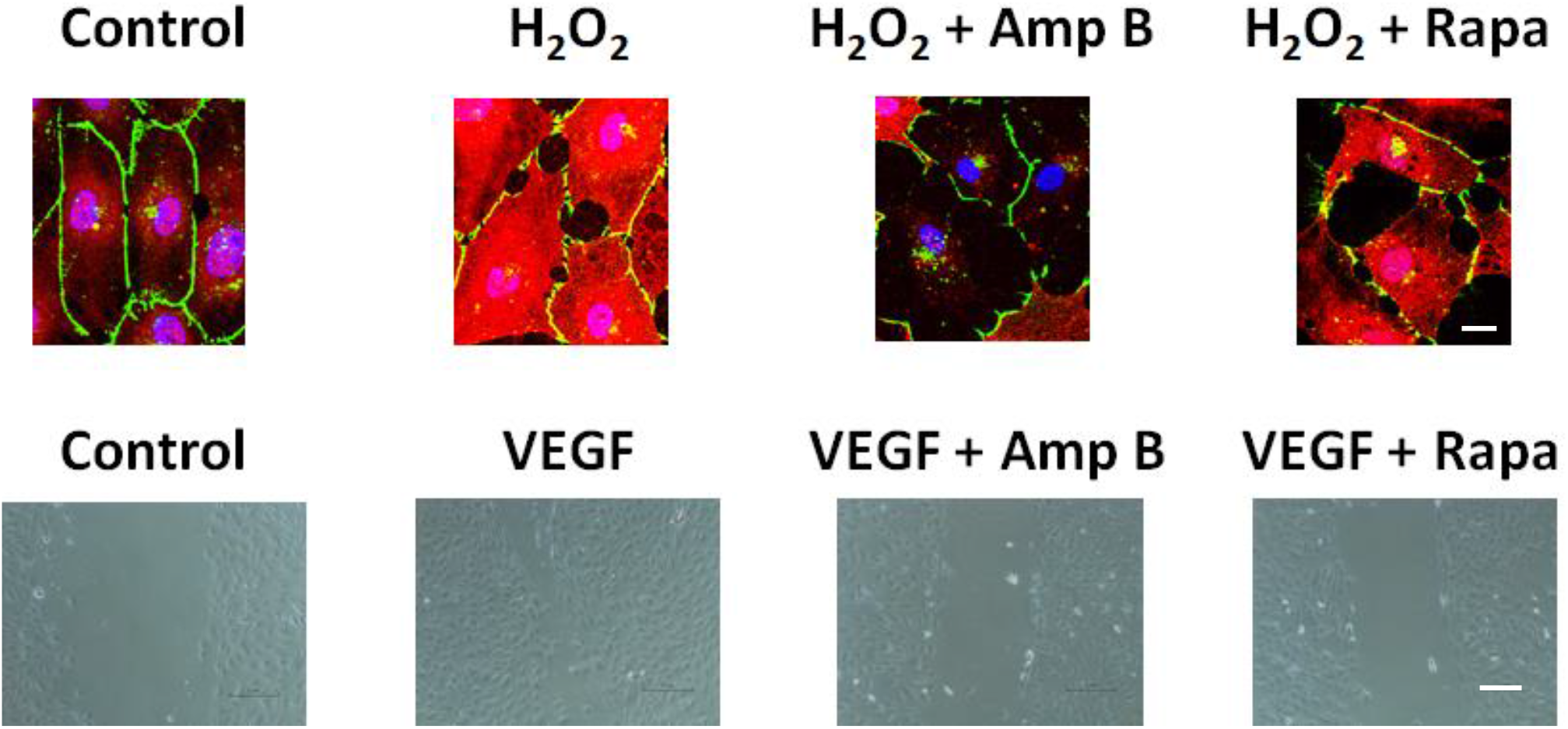
AMPhB and RAPA inhibits CLIC4 membrane translocation and endothelial cell migration. *Top Panel*: HUVECs (Control) immunostained for CLIC4 (red), VE-cadherin (cell junctions, green) and DAPI (nucleus, blue) under confluent conditions show the presence of CLIC4 within cytosol and nucleus with the maintenance of endothelial cell barrier indicated by the tight cell-cell junctions. HUVECs treated with 0.003% hydrogen peroxide (H_2_O_2_) significantly increased the expression levels of CLIC4, including at the plasma membrane with disruption to the barrier functions. These effects were partially reversed when treated with 10 nM AMPhB or 10 nM RAPA as indicated by decreased CLIC4 staining in the cells, especially at the plasma membrane. Scale bar = 50 μM **Bottom Panel:** HUVECs were seeded in 6-well plates and “scratch-wounded” using a universal 10 μl pipette tip. After pre-treated with VEGF (2ng/mL) for 2 h, cells were treated with 10 μM AMPhB or RAPA. Representative images of control cells and cells treated with VEGF with or without AMPhB or RAPA indicate inhibitory effects of these drugs on VEGF-induced cell migration known to be mediated by CLIC4. Scale bar = 100 μM.

Cell migration is a key indicator of many biological processes including inflammation, angiogenesis and cancer progression ^51^. Here we used a widely known *in vitro* cell migration assay method (wound healing assay) to examine the effect of AMPhB and RAPA on HUVECs with VEGF as a positive control to promote cell migration. As shown in Figure 4 (bottom panel), the migration rate promoted by VEGF is attenuated by both AMPhB and RAPA, confirming its anticipated anti-migratory effects on endothelial cells.

### 4.5. Site-specific binding of AMPhB and RAPA induced characteristic changes in CLIC4 structure and dynamics

Conformational events associated with the targeted binding of these inhibitor compounds to the protein were evaluated using an all-atom MD simulation approach. This was important to understand structural changes with respect to AMPhB and RAPA binding, as validated *in vitro*.

Firstly, the stability of the whole protein systems across the entire simulation time-frames relative to changes in Cα atomistic motions were measured using RMSD. As estimated, unbound CLIC4 (APO) exhibited a very high degree of structural instability until ∼375ns time-frame where it attained convergence with lowered deviations (Figure 5). Relatively, RAPA and AMPhB notably lowered the RMSD by ∼2Å indicative of their stabilizing effects on the protein. Corresponding mean RMSD values are shown accordingly in Supplementary Table 2. To minimize the effects of structural disorderliness (entropy), particularly in the unbound protein, stable trajectories from the last 50ns (450ns – 500ns) were retrieved for all systems and used for subsequent global analyses. RMSD distribution violin plots were employed to measure variations in CLIC4 conformation in the presence and absence of the compounds.

**Figure 5.**
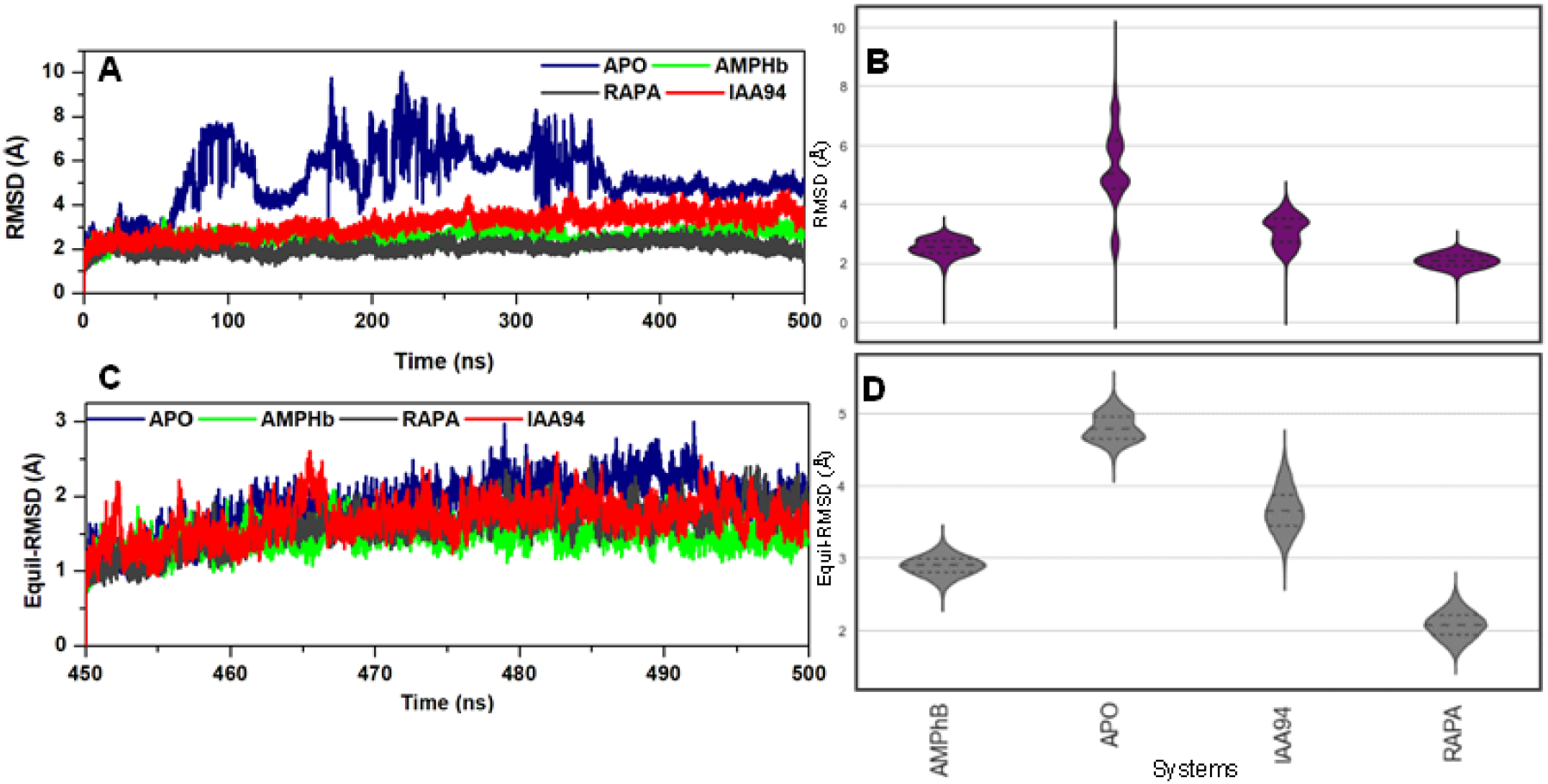
RMSD plots for unbound and ligand-bound CLIC4. [A] Whole time-frame (500ns) RMSD plot showing high deviations in unbound CLIC4 relative to stable ligand-bound systems. [B] RMSD distribution density plots for whole time RMSD. [C] Final equilibrated RMSD plots obtained from the terminal 50ns [D] RMSD distribution density plots for final equilibrated RMSD.

As shown in Figure 5D, unbound CLIC4 exhibited bimodal conformational distributions which indicates multiple conformations attained over the course of the simulation. Unimodal distributions were however observed in AMPhB-and RAPA-bound CLIC4 indicative of their roles in stabilizing the protein structure. The effects of the compounds on the compactness of the whole protein was further investigated using RoG calculations. From the plots (equilibrated timeframes), the binding of AMPhB and RAPA resulted in an increased protein RoG (_*AMPhB*_Mean_RoG (Å) = 19.83±0.12; _*RAPA*_Mean_RoG (Å) = 20.56±0.12) which correlates with the loss in structural compactness relative to the unbound protein (_*APO*_Mean_RoG (Å) = 19.36±0.10).

Furthermore, changes in residual fluctuations within the protein with respect to ligand binding were monitored using RMSF metrics and shown in Figure 6B. As observed, notable fluctuations occurred around residues 54-75 and 159-175 which, respectively, mapped out to the connecting loop and flexible foot loop regions of the protein. However, the intensity of fluctuation was highest in the presence of IAA-94. Peculiar to the binding of RAPA are fluctuations of residues 25-34, which constitute the catalytic (GSH-binding) loop while the binding of AMPhB specifically induced the flexibility of residues 80-84 that form an extended β-sheet loop from the GSH-binding site.

**Figure 6.**
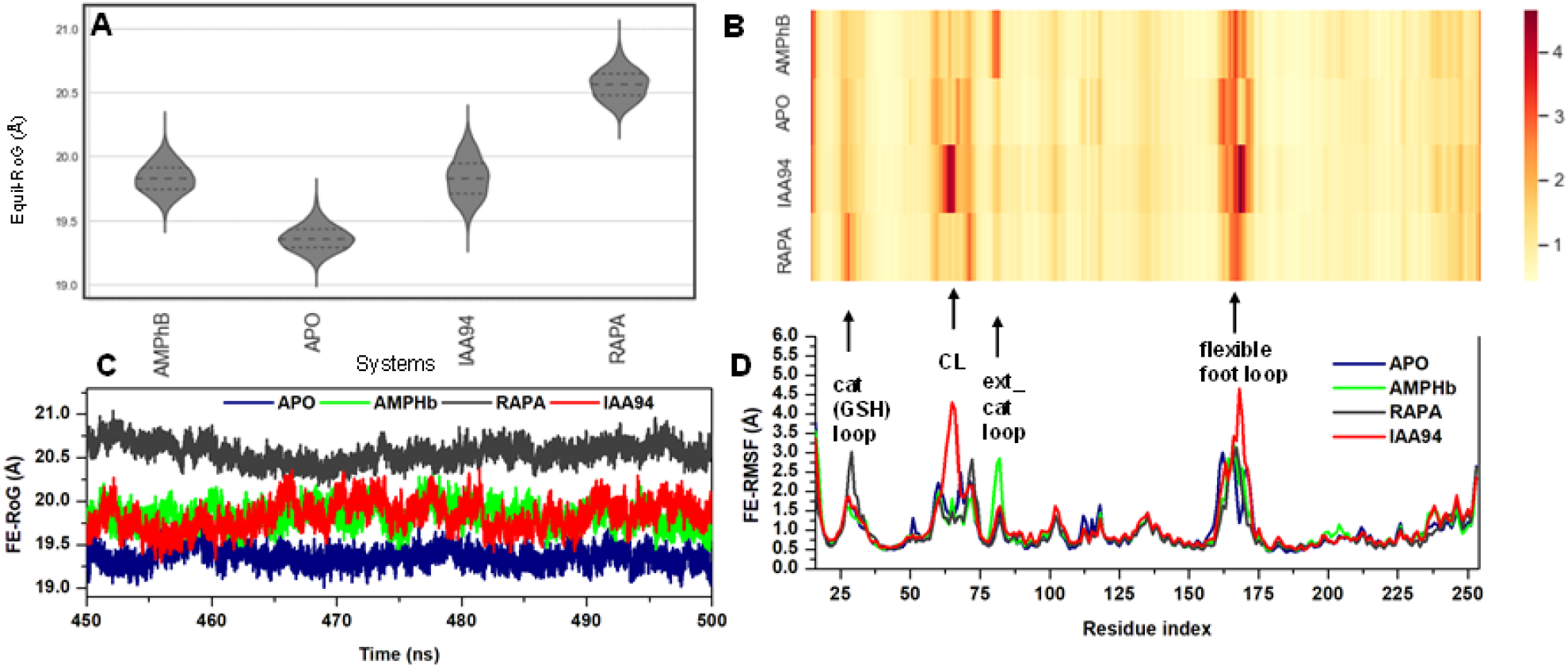
RoG and RMSF plots measuring variations in structural compactness and residual fluctuations [A] Density distribution plots of equilibrated RoGs [B] Heatmap showing fluctuations among of constituent residues [C] equilibrated RoG line plots [D] RMSF line plot mapping out corresponding residues and their degree of fluctuations as indicated in B.

Corresponding mean RMSF values for these structural elements are presented in Table 4. To corroborate these ligand-binding effects, the relative C-α stability and compactness were evaluated using the RMSD and RoG distributions (Figure 7A-E). Relative to other systems, RAPA allosteric induced a bimodal distribution at the catalytic (GSH) loop indicative of its distortive effect. More so, the binding of AMPhB caused a notable conformational alteration at the extended catalytic loop with a corresponding increase in Cα motions (Figure 7C). These loops distinctly impacted by RAPA and AMPhB are proximal to the GSH site and when distorted possibly interferes with the GSH-dependent activity of CLIC4 which is crucial to their cellular catalytic roles.

**Table 4:** Mean FE-RMSF calculations for important CLIC4 structural elements

| Residual fluctuation (Å) |  |  |  |  |
| --- | --- | --- | --- | --- |
| Structural elements | CLIC4 | CLIC4+control | CLIC4+AMPhB | CLIC4+RAPA |
| CL | 1.41±0.58 | 1.81±1.05 | 1.25±0.37 | 1.29±0.57 |
| Catalytic loop | 1.35±0.29 | 1.42±0.29 | 1.31±0.17 | 1.68±0.68 |
| FFL | 2.01±0.78 | 2.31±1.12 | 1.79±0.80 | 1.62±0.83 |
| N-to-C- loop | 0.89±0.30 | 0.99±0.30 | 0.88±0.31 | 0.87±0.24 |
| Ext_βcat_loop | 0.98±0.29 | 1.24±0.66 | 1.93 ±0.43 | 0.96±0.27 |

**Figure 7.**
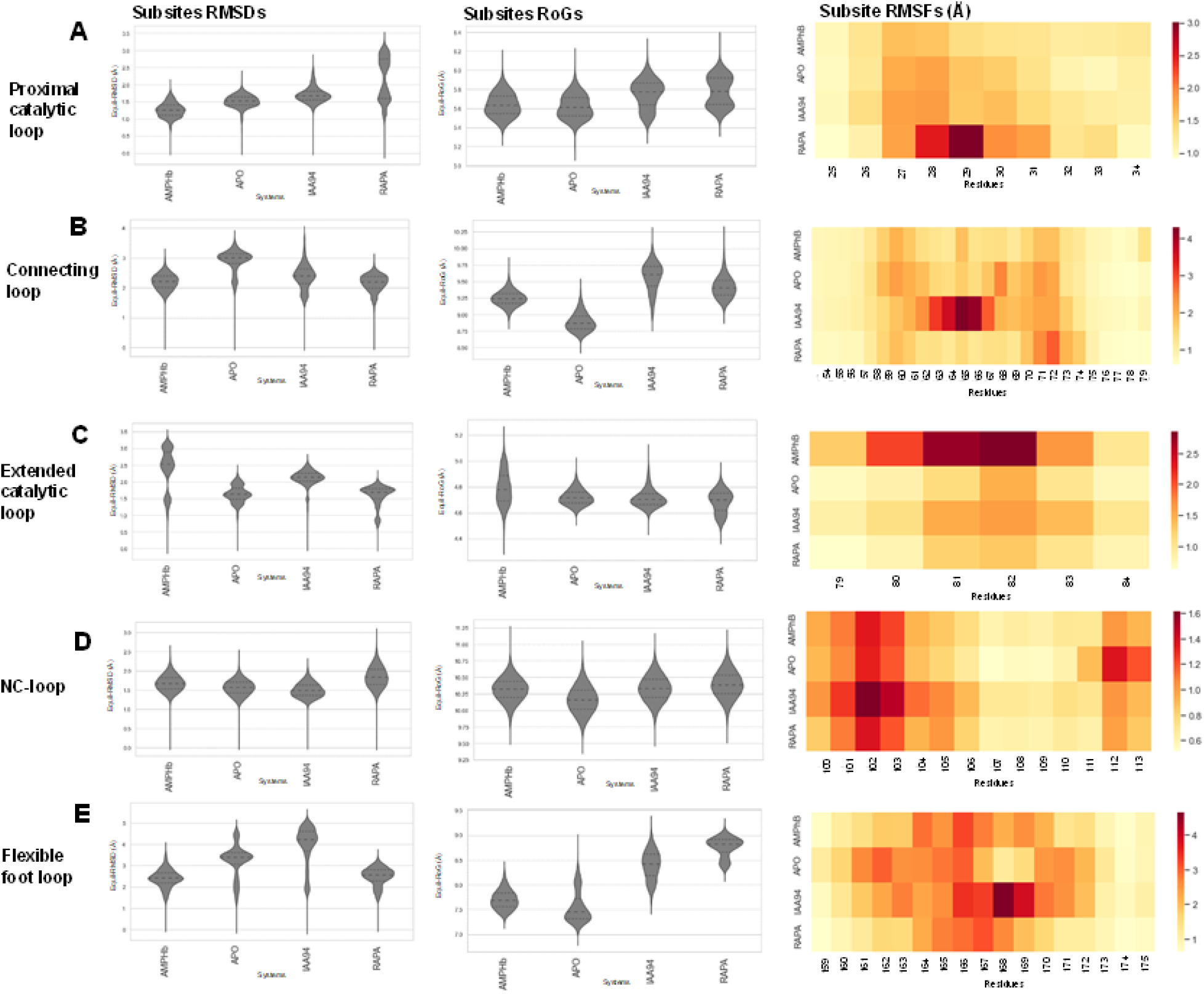
Conformational RMSD, RoG and RMSF analyses for important subsites on CLCI4 with notable ligand effects [A] proximal catalytic (GSH) loop [B] connecting loop [C] extended catalytic loop [D] N-to-C terminal connecting loop [E] flexible foot loop

This could underlie their inhibitory mechanisms. 3D representations of the degree of structural alteration at these regions are shown in Figure 8D-F. It is also important to mention that although none of the compounds elicited effects at the connecting N-C terminal loop, their effects were varied on the proximal flexible foot loop (Figure 7E). As seen, while high distortions characterized the loop in unbound and IAA-94 systems, it appeared to be more stable in CLIC4 bound by AMPhB and RAPA. This could as well impact on the mobility of the protein as this region is crucial for CLIC4 translocation from the cytoplasm to the membrane.

**Figure 8:**
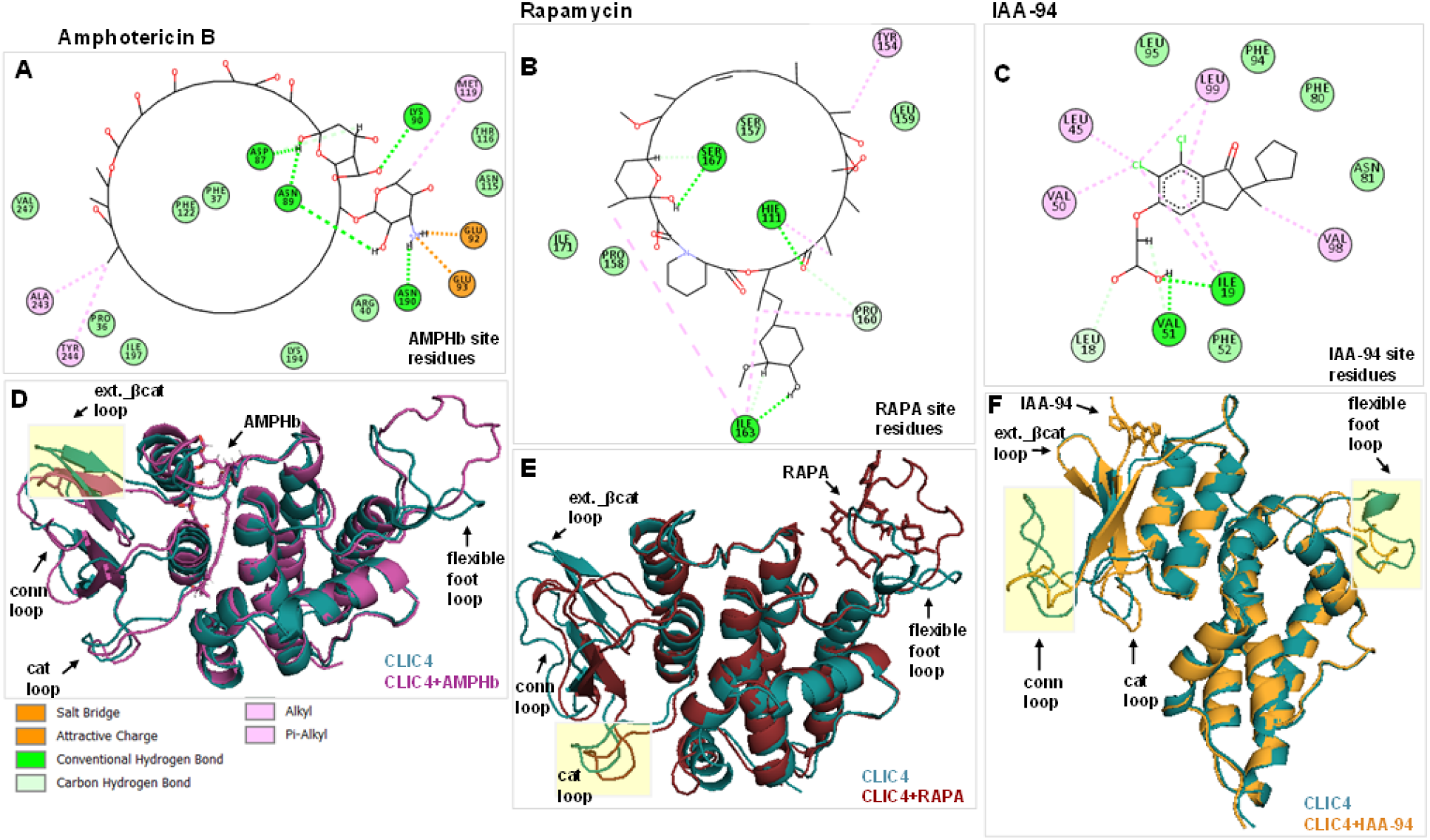
2D mapping of residue interactions for [A] CLIC4 and Amphotericin B [B] CLIC4 and Rapamycin [C] CLIC4 and IAA-94. 3D superposition of unbound and [D] Amphotericin B [B] Rapamycin [E] IAA-94-bound CLIC4 showing alterations at key regions.

Similarly, the connecting loop region exhibited a more stable conformations in the presence of AMPhB and RAPA but showed bimodal RMSD distributions in unbound and IAA94-bound CLIC4 (Figure 7B).

### 4.6. MM/GBSA calculations revealed mechanistic variations in the relative binding affinities of AMPhB and RAPA to CLIC4

The respective affinities (binding free energies, ***ΔG***_***bind***_) by which the inhibitors bind to CLIC4 were determined using the MM/GBSA method which also provided insights into the contributions of the various energy components to achieve ligands’ stable binding. This also was used as an approach to validate the initially derived docking affinities. It is important to further emphasize that the final equilibrated (stable) time-frames (last 50ns) were employed for the energy calculations in order to eliminate entropical effects (T*ΔS* = 0). From our calculations, AMPhB exhibited the most favorable ***ΔG***_***bind***_ of -41.36 kcal mol^-1^ while RAPA had a ***ΔG***_***bind***_ value of -23.08 kcal mol^-1^. Relatively, IAA94 had the least binding affinity with an energy value of -19.21 kcal mol^-1^ (Table 5). As observed, these findings also correlate with the order of the binding (Vina) scores.

**Table 5:**
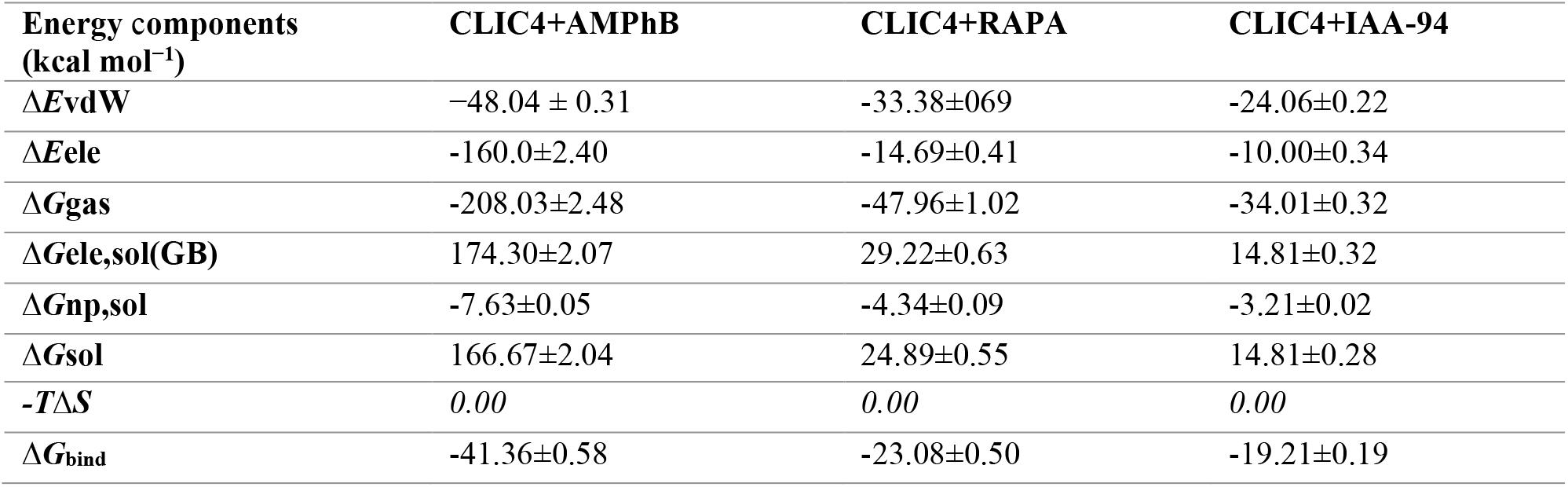
Binding free energy profiles of AMPHb, RAPA and IAA-94.

More so, electrostatic energies (**Δ*E*ele**) had the most contributions to the stable binding of AMPhB at Site 3 while van der Waals (**Δ*E*vdW**) was most prominent to RAPA and IAA-94 at their respective allosteric binding sites. Using the average structures, 2D mapping of the ligand-residue interactions further revealed the involvement of the terminal aminium group of AMPhB to form strong N=O (attractive charge-charge) bonds (*d* = 5.50Å) with E93 and a NH--O salt-salt bridge with E92 (*d* = 1.85Å) (Figure 8A). Also involved in interacting with the aminium group is N190 via conventional NH--O (*d* = 1.81Å) while the oxane moieties interacted via additional hydrogen (NH--O and CH—O) bonds with D87, N89, and K90. These strong interactions corroborate the high **Δ*E*ele** values estimated and the short bond distances (*d*) that ranged between 1.65 Å -5.5 Å further support the ligand-binding stability. Also, π (aromatic) interactions observed with M119, A243 and Y244 contribute towards the **Δ*E*vdW** which also impacts on ligand stability. A high Δ*G*ele,sol(GB) and Δ*G*sol for AMPhB indicates its interaction was more favorable within the Site 3 pocket and with constituent residues away from the regions accessible by solvent. This correlates with the estimated non-polar energy (Δ*G*np,sol) of -7.63 kcal mol^-1^ which favors the hydrophobic interaction of the inhibitor. Furthermore, π interactions were prominent in the binding of RAPA and involved Y154, P160, I163, H111 which could account for the high ***vdW*** energies. In addition, S167 interacted with the oxane group of RAPA while H111 formed conventional hydrogen (NH--O) with an extending carbonyl group. I163 also interacts with the 1-diol group of the terminal hexanol ring via a conventional NH--O altogether contributing to the binding stability of RAPA (Figure 8B). RAPA exhibited a favorable non-polar (hydrophobic) binding (Δ*G*np,sol) similar to IAA-94 which also had more ***vdW*** energies to attain a stable binding. Prominent to the binding of IAA-94 were π interactions involving I19, L45, V50, V98 and L99, while I19 and V51 contributed conventional H-bonds to IAA-94 binding (Figure 8C).

## 5. DISCUSSION

CLIC4 is not a typical ion channel protein but metamorphic in nature, which accounts for its involvement with numerous downstream pathways and effectors across diverse cell forms and processes. This functionality, however, makes it central to the development of various cancer and vascular diseases, among others. The significance of inhibiting CLIC4 to ameliorate pathologies have been previously reported and involved the use of research methods like gene knockout and RNAi approaches^18,52–54^. Notable is a study by Abdul Salam *et al*., which showed CLIC4/Arf6 inhibition and ameliorative effects in PAH ^18^. In spite, no specific CLIC4 inhibitor has been identified to date thereby limiting translatable interventions.

This study incorporates structure-based approaches and for the first time, reports small-molecule inhibitors of CLIC4. We also report novel allosteric (non-GSH) sites on the protein which are highly suitable for targeting by chemical compounds or entities. We also demonstrated, experimentally, the potentials of the identified hit compounds to specifically bind CLIC4 and as well reverse CLIC4-mediated oxidative stress in endothelial cells. Also, we experimented the anti-migratory effects of the compounds on endothelial cells.

The soluble form of CLIC4 is structurally related to omega-type glutathione -transferases (GST-omega) and thought to exhibit GSH-dependent enzymatic activity ^55^. This accounted for the use of IAA-94, a general chloride channel blocker, as the control compound in this study. Though its CLIC binding property has been widely reported, the specific binding region on CLIC4 remains elusive. Our inhibitor screening study demonstrated that it potentially binds around the GSH binding site which is most likely due to its chemical similarity with ethacrynic acid, a known GST inhibitor. However, its binding affinity is relatively low compared to the hit compounds and our experimental studies corroboratively revealed that IAA-94 poorly binds to CLIC4, relative to the hit compounds. From the NMR data, IAA-94 failed to induce any significant chemical shift perturbations on the NMR spectrum of CLIC4, indicating its inability to bind CLIC4 in this soluble form. This could also indicate that conformational changes induced by IAA94 as observed in the MD studies have no translatable effects *in vitro* and that many of the inhibitory effects widely reported in literature may be due to indirect effects or limited to the channel form alone

The lack of information on possible allosteric (non-GSH) and druggable sites on CLIC4 has not favored previous implementation of structure-based discovery of potential CLIC4 inhibitors. This further explains the significance of this present study and how it aids future research efforts. Identifying allosteric inhibitors is more advantageous in the drug development process as it provides a more feasible avenue to discover drug molecules with high target specificity. This is because, contrary to orthosteric sites, allosteric sites are less conserved, hence drugs binding to these regions are more specific and most likely, less toxic to human^56,57^.

Importantly, we identified two high-affinity sites in CLIC4 other than the known GST-like site that is highly conserved among the CLIC protein family. An interesting finding was the identification of the flexible foot loop region (Site 2) and its high potentials for allosteric targeting by small-molecule compounds. Consequentially, a large proportion of the predicted hit compounds interacted preferentially at this region with high affinities. Functionally, the flexible foot loop region is crucial for the membrane translocation of CLIC proteins, and, if effectively inhibited, could prevent CLIC4 cellular motility which is essential for various pathological involvement. This study therefore opens up avenues to explore site targetability, particularly, the identification of crucial interactive residues such as P158, L159, Thr175 and Arg176 among others, which will be essential for future site-specific structure-based inhibitor design studies.

Although our predicted inhibitors (amphotericin B and rapamycin) did not significantly impact on the flexible foot loop region, our MD simulation study revealed their respective allosteric effects were more prominent on the catalytic loops. Distortions in key catalytic region of CLIC4 as induced by these proteins could in turn affect its enzymatic activity and to a larger extent, effector protein interactions. Corroboratively, NMR results revealed that both rapamycin and amphotericin B induced structural changes in CLIC4. While the low water solubility of rapamycin makes it impossible to compare affinities, both molecules display clear chemical shift perturbations on a small subset of peaks in the ^1^H-^15^N NMR spectrum of CLIC4. Additionally, residues involved in high-affinity interactions with inhibitor molecules are essential for binding stability and such residues; Asp87, Asn89, Lys90, Glu92, Glu93 and Asn190 for amphotericin B, and His111, Pro160, Ile163, and Ser167 for rapamycin could be explored in future studies for discovering ligands with improved specificity for both sites. Furthermore, the impact of both compounds on CLIC4 enzymatic activity could correlate with their abilities to ameliorate CLIC4-induced cellular stress and reverse VEGF-mediated cell migration. *In vitro* validation assay using endothelial cell system further reflects the functional effects of the NMR and MD binding observations and showed that the tested compounds were able to inhibit CLIC4-mediated endothelial response especially in perpetrated pathological conditions. The precise mechanism of this however, needs further investigation but is likely to involve the VEGF or SIP-1 pathway.

In summary, we employed structure-based methods that led to the identification of amphotericin B and rapamycin as allosteric inhibitors of CLIC4. Experimental validation studies further confirmed their binding potentials and ability to reverse CLIC4-mediated cellular dysfunctions. This presents an important advancement in therapeutic strategies to specifically target the pathological involvement of CLIC proteins.

## Acknowledgement

We would like to thank the Center for High-Performance Computing, Cape Town, South Africa for providing computational resources.

## SUPPLEMENTARY INFORMATION

**Supplementary Table 1:**
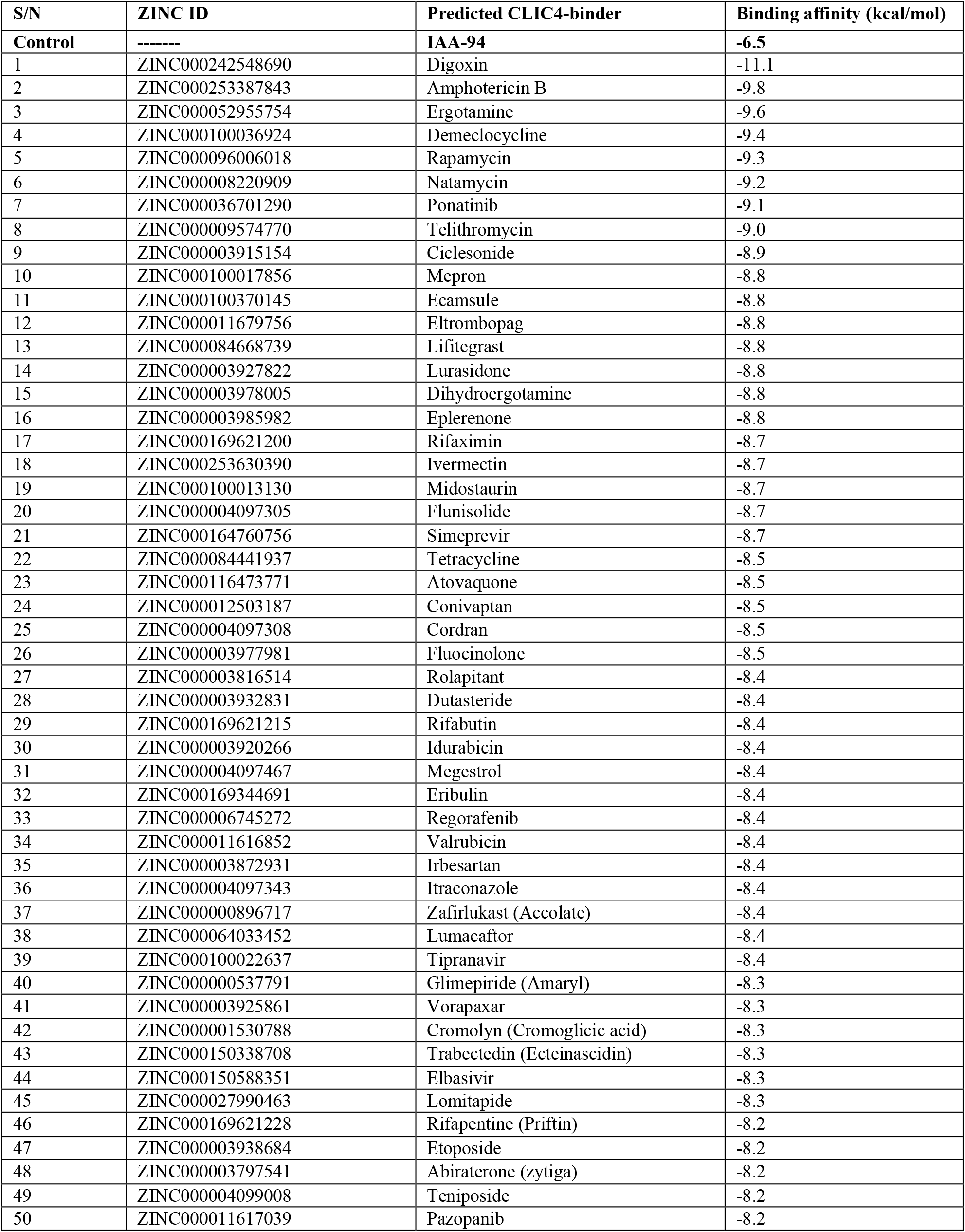
Corresponding docking scores of the top 50 potential CLIC4 inhibitors and control compound, IAA94

**Supplementary Table 2:** Estimations of differential conformational changes in CLIC4

| **Structural analysis (Å)** | **CLIC4** | **CLIC4+IAA94** | **CLIC4+AMPHb** | **CLIC4+RAPA** |
| --- | --- | --- | --- | --- |
| Whole RMSD | 5.22±1.40 | 3.12±0.53 | 2.53±0.31 | 2.07±0.24 |
| FE-RMSD | 1.90±0.34 | 1.69±0.26 | 1.45±0.18 | 1.61±0.28 |
| FE-RoG | 19.36±0.10 | 19.79±0.16 | 19.83±0.12 | 20.56±0.12 |

